# Role of duplicate genes in determining the tissue-selectivity of hereditary diseases

**DOI:** 10.1101/171090

**Authors:** Ruth Barshir, Idan Hekselman, Netta Shemesh, Moran Sharon, Lena Novack, Esti Yeger-Lotem

## Abstract

A longstanding puzzle in human genetics is what limits the clinical manifestation of hundreds of hereditary diseases to certain tissues or cell types, while their causal genes are present and expressed throughout the human body. Here we considered a possible role for paralogs of causal genes in affecting this tissue selectivity. It has been shown across organisms that paralogs can compensate for the loss of each other. We hypothesized that specifically in the disease tissue causal genes and their paralogs are imbalanced, leading to insufficient compensation and to the emergence of disease phenotypes. While demonstrated previously in the context of few specific diseases, this hypothesis was never assessed quantitatively at large-scale. For this, we analyzed functional relationships between causal genes and their paralogs associated with 112 tissue-selective hereditary diseases. To test our hypothesis we used several large-scale omics datasets, including RNA sequencing profiles of over 30 different human tissues. Indeed, the expression of causal genes and their paralogs was significantly imbalanced in their disease tissues compared to unaffected tissues. Imbalanced expression was evident across different disease tissues, and was common to causal genes with single or multiple paralogs. This imbalance was driven by significant upregulation of the causal gene in its disease tissue, often combined with significant down-regulation of a paralog. Nevertheless, in additional 20% of the causal genes, a paralog alone was significantly down-regulated in the disease tissue. Our results suggest that dosage relationships between paralogs affect the phenotypic outcome of germline aberrations, adding paralogs as important modifiers of disease manifestation.

## INTRODUCTION

Hereditary diseases are caused by germline aberrations that are common to cells throughout the human body. For hundreds of these diseases, these germline aberrations have been identified and mapped to causal genes [1], and many more causal genes are likely to be identified in coming years owing to the extensive usage of sequencing techniques in medical settings [2]. However, the identification of a causal gene is often just the starting point for understanding the molecular basis of each disease. The genotype-to-phenotype relationship between a causal gene and the respective disease phenotype is typically complex [3,4], and, for numerous hereditary diseases remains to be elucidated. By shedding light on these relationships, we hope to obtain better understanding of disease mechanisms and advance the search for cures.

Tissue-selectivity is a hallmark of many hereditary diseases [5]. For example, familial mutations in BRCA1 gene increase the risk for breast and ovarian cancers, and familial mutations in RB1 gene lead primarily to retinoblastoma. From an evolutionary point of view, tissue-selectivity is not surprising given that limited manifestation is probably less detrimental than whole body diseases, and thus more likely heritable. Yet, tissue-selectivity is intriguing due to the pattern of expression of causal genes. For example, both BRCA1 and RB1 are expressed ubiquitously across most human tissues without eliciting disease phenotypes in those tissues. In fact, most causal genes exhibit tissue-specific disease manifestation along with tissue-wide expression [5,6]. Several molecular mechanisms may lead to this phenomenon. In some cases, the disease-manifesting tissue (denoted disease tissue henceforth) has unique features [7,8], such as long-lived neurons and age-related protein misfolding diseases [9]. In other cases, the tissue-selective effect may depend on the specific isoform expressed in that tissue [10]. Meta-analysis studies showed that causal genes tend to have elevated expression preferentially in their disease tissues [5,6], hinting to a quantitative basis for tissue selectivity. Previously, we showed that causal genes tend to form tissue-specific interactions preferentially in their respective disease tissues, suggesting that these interactions contribute to tissue selectivity [5]. Yet for many hereditary diseases, the molecular mechanisms that underlie them remain hidden.

Here, we consider the role that paralogs of causal genes may play in determining the tissue-selectivity of hereditary diseases. Paralogs, namely homologous genes within the same species resulting from gene duplication events, have been repeatedly shown to have redundant functions and to compensate for the loss of each other (reviewed in [11]). At a systems level, paralogs were shown to be less essential than genes lacking paralogs (singletons) in yeast [12], worms [13], mice [14] and plant [15]. A recent measurement of the essentiality of over 17,000 human genes showed that the same tendency holds for human paralogs [16]. The impact of paralogs was also demonstrated in the context of disease. For example, a mouse model of retinoblastoma that carries a homozygous deletion in the Rb gene, the homolog of human RB1 gene, does not develop retinoblastoma [17], unless one of the paralogs of Rb, p107 [18] or p130 [19], is removed.

Interestingly, in several cases the compensatory impact of paralogs was found to be dosage-dependent. For example, mouse embryos that are homozygous for Mek1 gene deletion and which typically die due to placental defects, survive if two copies of Mek2 gene are inserted, while one copy of Mek2 is not sufficient [20]. Similarly, the Eif2s3y gene on the mouse Y chromosome was shown to be replaceable by its X-linked homolog Eif2s3x gene for spermatogenesis initiation, but more copies of Eif2s3x were required for progression through meiosis [21]. In human, the essentiality of each of the two paralogous helicases, the genes DDX3Y and DDX3X, was inversely correlated with the expression level of the other paralog, stressing that their functional redundancy is dosage-dependent [16].

We hypothesized that tissue-selectivity of some hereditary diseases may be related to quantitative relationship between causal genes and their paralogs. Accordingly, owing to the functional redundancy between paralogs, a paralog of an aberrant causal gene can generally compensate for its malfunction (Figure 1A). However, when the quantitative relationships between them change, compensation may become insufficient and disease phenotypes will emerge. This might occur when the causal gene is up-regulated in the disease tissue without a similar change in the level of the paralog (Figure 1B). Alternatively, the causal gene may be expressed at an intermediate level in the disease tissue, but the paralog is down-regulated at the disease tissue (Figure 1C). While dosage-dependent compensation relationships between paralogs were demonstrated previously (e.g., [16,20,21]), this phenomenon was never analyzed systematically at large-scale in the context of tissue-selective diseases.

**Figure 1.**
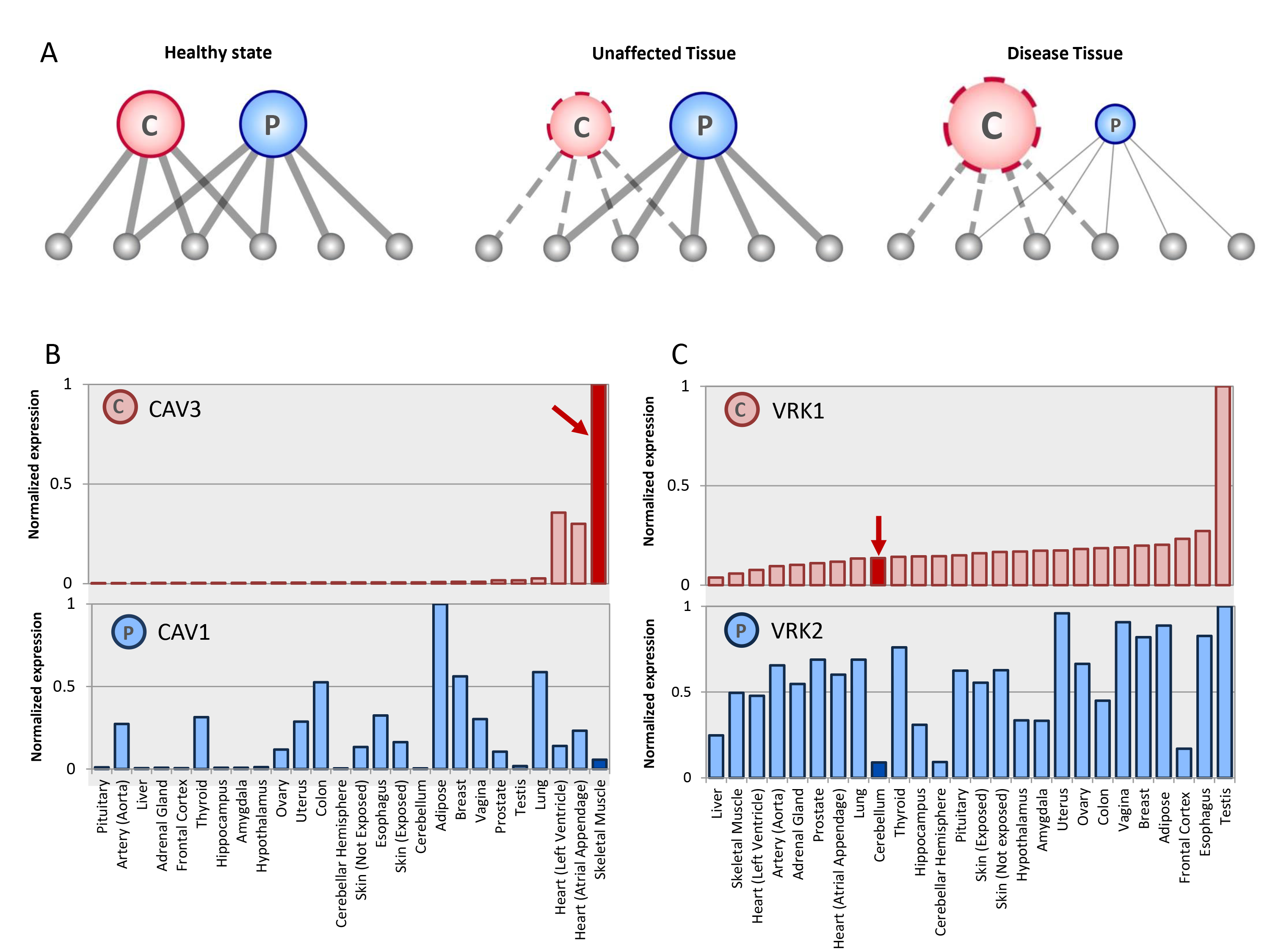
The functional redundancy between causal genes and their paralogs is dosage-sensitive. A. A network model representing the functional redundancy between a causal gene and its paralog. In the healthy state (left), the causal gene (marked C) and its paralog (marked P) have redundant functions, represented as common interactors. In the aberrant state, the casual gene has limited functionality (dashed edges). In unaffected tissues (middle), the limited functionality is masked by the presence of its paralog. In the disease tissue (right), masking is reduced (thin edges) due to relatively low expression of the paralog, and the limited functionality is exposed. B. Limited masking in the disease tissue caused by over-expression of the causal gene. Germline mutations in CAV3 lead to muscular dystrophy. In muscle (red arrow) CAV3 is expressed at its highest level whereas its paralog, CAV1, is relatively lowly expressed. C. Limited masking in a disease tissue may arise from under-expression of the paralog. Germline mutations in VRK1 cause pontocerebellar hypoplasia. In the disease tissue (cerebellum, red arrow), VRK1 is expressed at an intermediate level, whereas its paralog, VRK2, is significantly under-expressed. Expression data were downloaded from GTEx portal [40] (see Methods). Gene expression levels were normalized to their maximum levels across 45 tissues (26 tissues shown). Expression in disease tissues appears as bold color bars.

In this study, we analyze quantitatively the relationships between 81 causal genes and their paralogs across 112 hereditary diseases. We focused on hereditary diseases that manifest selectively in distinct tissues, including the brain, skeletal muscle, heart, skin, liver, thyroid or testis. We took advantage of various omics data including 421 RNA-sequencing profiles of human tissues made available by the Genotype-Tissue Expression (GTEx) consortium [22], to assess quantitatively the relationships between causal genes and their paralogs across 45 tissues. The majority of the causal genes were functionally-overlapping with their paralogs, and causal genes were less essential than singleton genes. Next, we computed the expression ratios between causal genes and their paralogs in different tissues. The ratios were typically similar across tissues, except for the disease tissues where the ratios were significantly high. These high ratios were observed for causal genes with different numbers of paralogs and in various disease tissues. To distinguish between the possible scenarios leading to imbalanced expression (Figure 1B,C), we carried differential expression analysis across the different tissues. In 27% of the cases, the causal gene was significantly up-regulated in the disease tissue. In 20% of the cases a paralog was significantly down-regulated in the disease tissue, and in additional 22% of the cases both occurred. These results suggest that paralogous compensation can shed light on the tissue-selective manifestations of hereditary diseases.

## RESULTS

### Functional relationships between causal genes and paralogs

We started by creating a high-confidence dataset of tissue-selective hereditary diseases. For this, we manually curated hereditary diseases that manifested clinically in one of the following tissues: brain, skeletal muscle, heart, skin, liver, thyroid or testis. We included diseases with various modes of inheritance, since in both dominant and recessive disorders paralogous compensation may contribute to the robustness of unaffected tissues. For each disease, we extracted its known causal on phylogeny and sequence identity (see Methods). The causal genes contained various types of aberrations, several of which can lead to partial or complete loss of the gene product and its function, due to, e.g., protein truncation or in-frame missense mutations [23,24]. Other aberrations could lead to gain-of-function for which paralogous compensation may not be relevant, but these were shown in a systematic screen to occur at low frequency [23]. Next, we examined the pattern of expression of causal genes across tissues, to avoid causal genes that are tissue-specific and thus inevitably elicit tissue-specific phenotypes. For this, we used RNA-sequencing profiles of human tissues made public by the GTEx consortium [22] (see Methods). We computed the number of tissues expressing each causal gene above a certain threshold (see Methods), and limited our analysis to causal gene and their paralogs that were co-expressed in at least five tissues.

Henceforth, we analyzed 112 hereditary diseases caused by germline mutations in 81 causal genes (Figure 2A and Table S1). The majority of the causal genes were expressed ubiquitously across tissues (83%, Figure 2B and Figure S1). Thus, in agreement with previous studies [5,6], tissue-selectivity of their respective diseases could not stem simply from tissue-selective expression of the casual genes. Most causal genes were associated with one or two paralogs (78%, Figure 2C), and many paralogs were also globally expressed (66%, Figure 2B).

**Figure 2.**
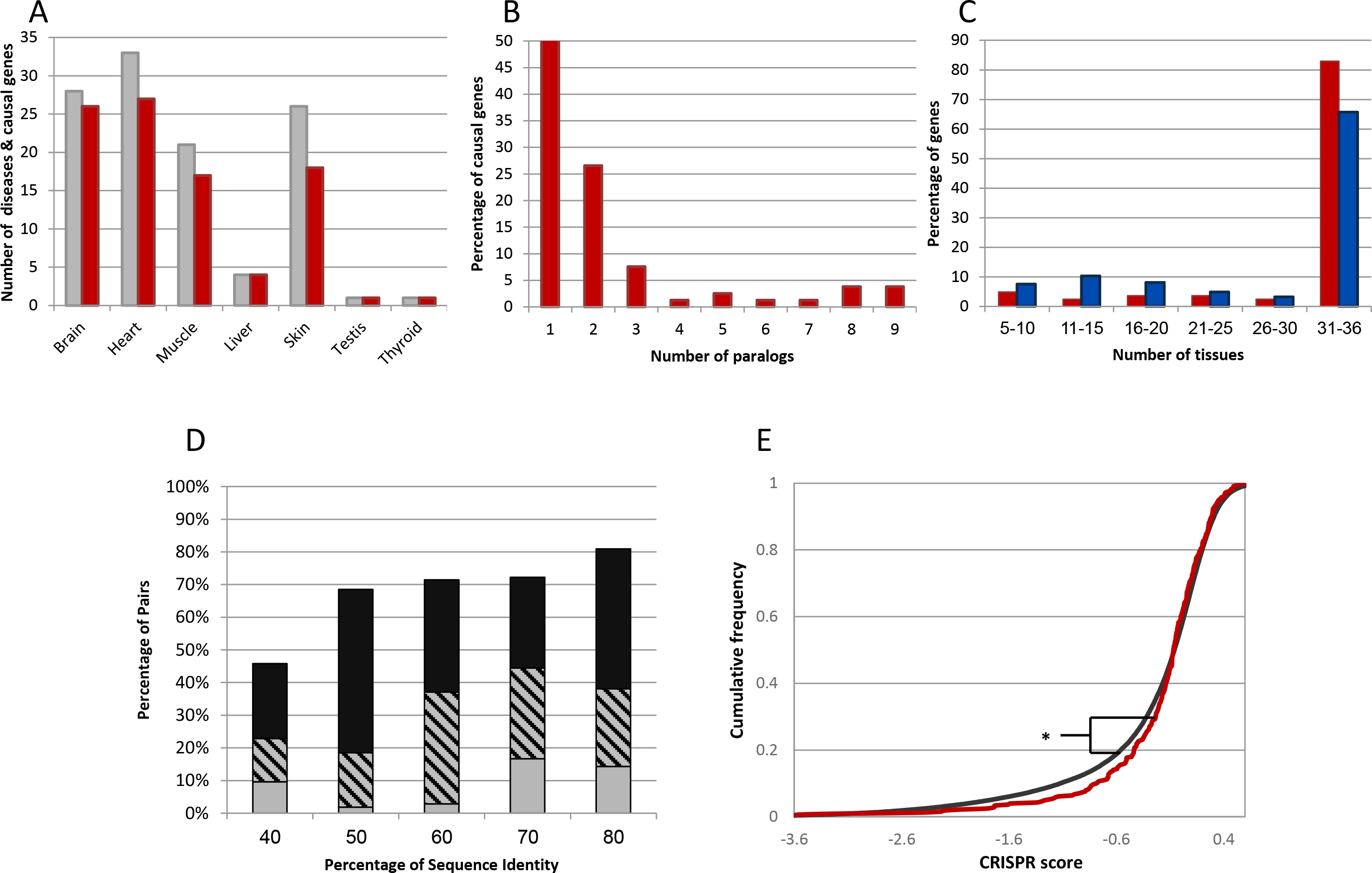
Evidence for functional overlap between causal genes and their paralogs. A. An overview of the manually-curated dataset of 112 tissue-selective hereditary diseases (gray bars) and their causal genes (red bars) by their disease tissues. B. The distributions of causal genes (red bars) and paralogs (blue bars) by number of expressing tissues, showing that most genes were expressed in all tissues. C. The distribution of causal genes according to the number of their paralogs. Most causal genes have up to two paralogs. D. The fraction of functionally-overlapping pairs among causal-gene and paralog (CGP) pairs with similar sequence identity levels. Evidence for functional overlap included significant co-expression across tissues (Pearson correlation p<0.01, gray bars), significant overlap in physical interaction partners (Fisher exact test p<0.01, black bars), or both (striped bars). The fraction of functionally redundant pairs increased with their sequence identity. E. The cumulative distributions of essential genes among causal genes (red) and protein-coding singleton genes (black) show that the causal genes in our dataset are significantly less essential than singleton genes (Kolmogorov-Smirnov test, p=0.02). The X axis shows cellular growth in the presence of inactivating mutations, denoted CRISPR score, where negative values mark essential genes [16].

Monogenic disease genes and their paralogs were shown previously to be frequently functionally overlapping, based on their co-expression relationships [25] and overlap in interaction partners [26]. We used similar measures to test for functional overlap between causal genes and their paralogs in our dataset. For each causal gene and its paralog, denoted causal gene – paralog (CGP) pair, we computed the expression correlation between them across all tissues, and the overlap in their protein interaction partners (see Methods). The majority of the CGP pairs were significantly functionally overlapping by at least one measure, and, as expected, functional overlap was more frequent among pairs with higher sequence identity (Figure 2D).

The functional overlap between causal genes and their paralogs suggests that causal genes in our dataset would have a lower tendency to be essential, relative to singleton genes. This was recently shown for human genes with paralogs in general [16]. We repeated the same test for the causal genes in our dataset (Fig 2E). Indeed, causal genes were significantly less essential than singleton genes (Kolmogorov-Smirnov test, p=0.012), in agreement with the presence of a functionally redundant paralog.

### Imbalanced expression of causal genes and paralogs occurs preferentially in the disease tissue

According to the imbalance hypothesis, the relative levels of a causal gene and its paralog are comparable across tissues, except for the disease tissue, where we expect to find a shift in balance (Figure 1). To test this hypothesis, we computed the ratio between the expression levels of causal genes and their paralogs across the different tissues (see Methods). We then compared the ratios obtained in disease tissues to the ratios obtained in other tissues (Figure 3A, left). The ratios in the disease tissue were significantly higher than the median ratios in unaffected tissues, in accordance with the imbalance hypothesis (Mann-Whitney, p<10^-15^). A similar shift in balance was evident upon considering only causal genes with a single paralog (Figure 3A, middle, p=0.0058). For causal genes with multiple paralogs, we additionally tested whether imbalance was still observable upon combining all the paralogs of each causal gene (see Methods). Indeed, these ratios too were significantly higher in the disease tissue relative to unaffected tissues (Figure 3A, right, p=0.001).

**Figure 3.**
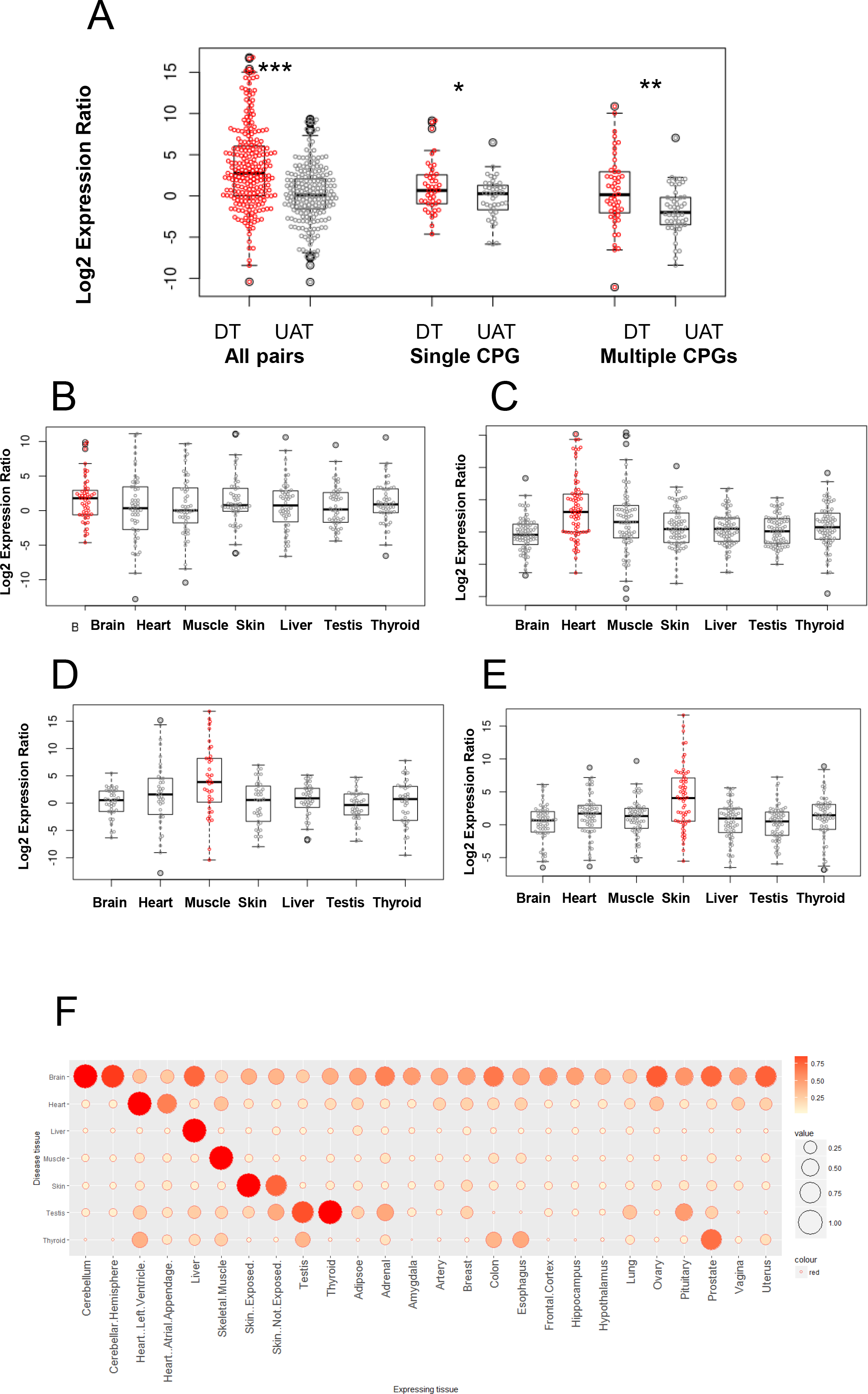
The expression of causal genes and their paralogs is highly imbalanced in their disease tissues compared to unaffected tissues. A. The ratio between the expression levels of causal genes and their paralogs across tissues. Each point represents a single pair; the ratio in the disease tissue appears in red, the median ratio in unaffected tissues appears in gray. Ratios in the disease tissue (DT) were significantly higher than in unaffected tissues (UAT) when all pairs were considered (‘All pairs’, Mann-Whitney test, p<10^-15^); when only causal genes with single paralogs were considered (‘Single CGP’, Mann-Whitney test, P=5.8*10^-3^); and when causal genes with multiple paralogs were considered and ratio was computed against their combined expression (‘Multiple CGP’, Mann-Whitney test, P=1.2*10^-3^). B. – E. The ratios between the expression levels of causal genes and their paralogs shown separately for causal genes sharing the same disease tissue. Each point represents the ratio observed in the disease tissue (red) and in an unaffected tissue (gray). The panels show genes causal for diseases manifesting in the brain (B), heart (C), skin (D), and skeletal muscle (E). In all panels, the median ratio is highest for pairs in their respective disease tissues. Additional disease tissues appear in Figure S2. F. The median ratios between the expression levels of causal genes and their paralogs across tissues. Each row corresponds to genes causal for diseases that manifest in the tissue designated on the left, and represents the median ratios per tissue normalized to the maximum in that row. In each row the median ratios in the disease tissue are highest. For brevity, 26 out of 45 tissues were shown.

We further tested the generality of the imbalance by dividing causal genes according to their disease tissues and repeating this test. Notably, the ratios obtained for pairs in their respective disease tissue were higher than the ratios obtained for the same pairs in the six unaffected tissues, for all disease tissues except testis and thyroid, which included a single causal gene (Figure 3B–E and Figure S2). We extended this test to include all tissues. Upon comparing the ratios for CGP pairs in their disease tissue to their ratios in all other tissues, we find that with the exception of testis and thyroid, the highest ratios were obtained consistently in the disease tissue and its closely related tissues, such as different regions of the heart, skin that is sun-exposed and unexposed, and closely related brain regions (Figure 3F and Figure S3). Thus, imbalanced expression of causal genes and their paralogs is prevailing among hereditary diseases.

### Down-regulation of paralogs can contribute to expression imbalance

Our next goal was to analyze quantitatively the causes for the shift in balance observed for causal genes and their paralogs in their respective disease tissues. In general, their balance could be shifted due to up-regulation of the causal gene, or down-regulation of its paralog, or both, as shown schematically in Figure 4A. The CGP pair CAV1, CAV3 presented in Figure 1B demonstrates the first scenario, while the CGP pair VRK1, VRK2 presented in Figure 1C demonstrates the second scenario. To distinguish between these scenarios and to quantify their frequency in our dataset, we used differential expression analysis. We focused on the 36 tissues for which five or more samples were available. For each disease tissue, we calculated the differential expression of genes in this tissue relative to all other tissues. This allowed us to identify rigorously genes that were up-regulated or down-regulated significantly in the respective disease tissue (2-fold change and p<0.01, see Methods). We then analyzed the frequency of the different scenarios among our CGP pairs (Figure 4B).

**Figure 4.**
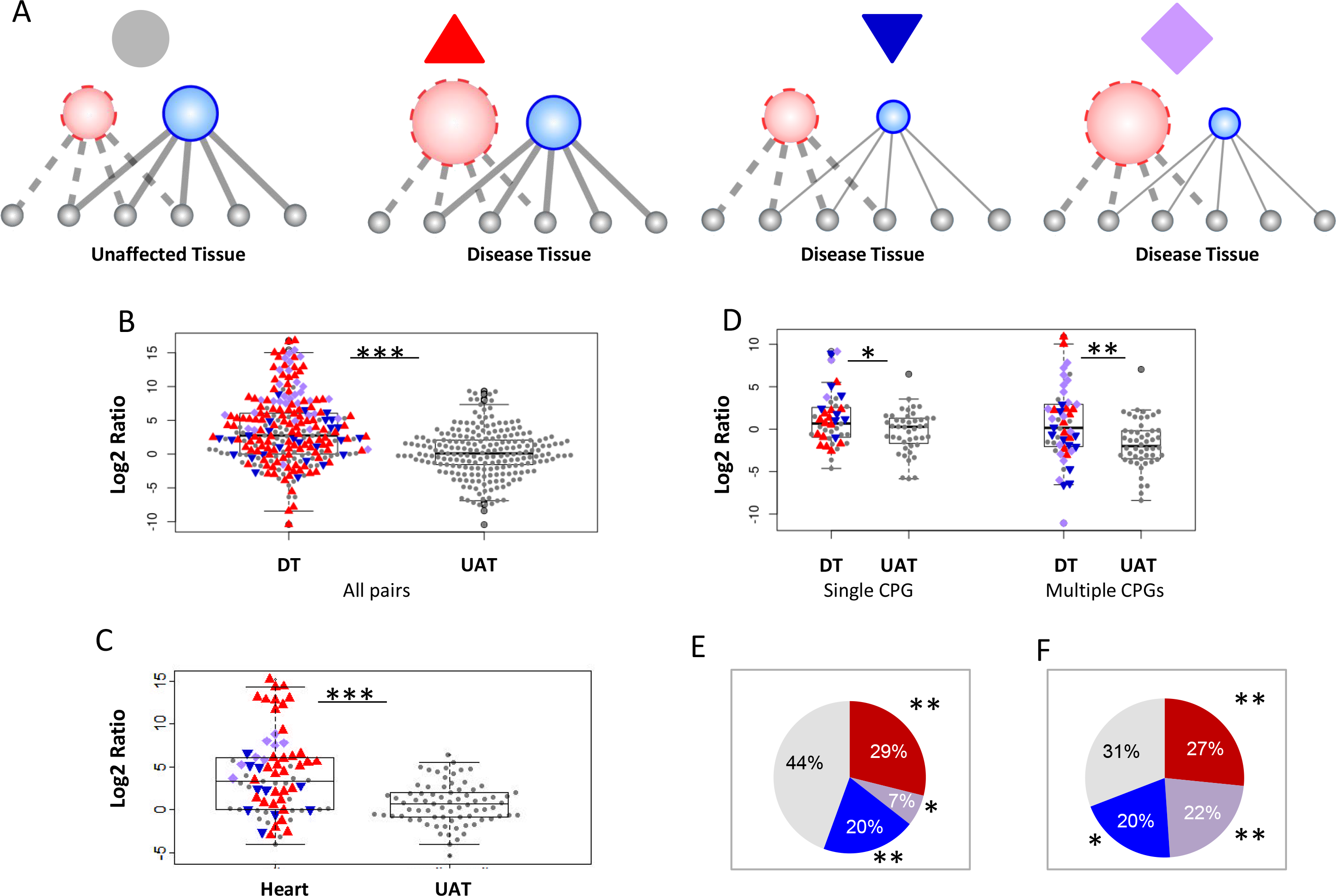
Differential expression analysis reveals causes for imbalanced expression of causal genes and paralogs in the disease tissue. A. Scenarios leading to imbalanced expression in the disease tissue. From left to right: a reference state; significant up-regulation of the causal gene; significant down-regulation of the paralog; significant up-and down-regulation of the causal gene and its paralog, respectively. B. The ratio between the expression levels of causal genes and their paralogs visualized according to the differential expression of the causal gene and its paralog. Each point represents the ratio of a specific pair in their disease tissue (DT) and unaffected tissues (UAT). Colors indicate whether in the disease tissue the causal gene was significantly over-expressed (red), the paralog was significantly under-expressed (blue), both co-occurred (purple), or none occurred (gray). In most pairs at least one pair mate was differentially expressed in the disease tissue. C. A visualization of the ratios between causal genes and their paralogs for causal genes with a single paralog (single CGP) or multiple paralogs (multiple CGP). For multiple CGPs, the ratio was computed against the combined expression of paralogs. Blue indicates that at least one paralog was significantly under expressed. D. The ratios between causal genes and their paralogs for genes causal for heart diseases. Data for other disease tissues appear in Figure S3. E. The frequency of imbalanced scenarios among the 45 causal genes with a single paralog. 29% of the causal genes were up-regulated in their disease tissues (red, p<10^-3^), 20% of the paralogs were down-regulated in the disease tissue (blue, p<10^-3^), and 7% of the causal genes had both (purple, p=0.05). F. The frequency of imbalanced scenarios among the 94 causal genes in our dataset. 26% of the causal genes were up-regulated in their disease tissues (red, p<10^-3^), 19% of the causal genes had a down-regulated paralog in the disease tissue (blue, p=0.024), and 24% of the causal genes had both (purple, p<10^-3^).

The most common scenario was the up-regulation of the causal gene, which we observed in 52% of the CGP pairs. In additional 15% of the pairs, this was combined with down-regulation of the paralog. Notably, in another 9% of the pairs the paralog alone was significantly down-regulated. To test the generality of these trends, we repeated this analysis for several partitions of the causal genes. We observed the same trends upon analyzing separately genes that share the same disease tissue (e.g., Figure 4C and Figure S4). The different scenarios were evident also when analyzing causal genes with a single paralog (Figure 4D). Specifically, 36% of the causal genes were up-regulated, including 7% where the paralog was down-regulated. In additional 20% of the genes, only the paralog was down-regulated (Figure 4E). We further tested whether these scenarios occurred preferentially in the disease tissue by carrying randomization tests (see Methods). We found that each scenario was significantly more frequent in the disease tissue than in unaffected tissues. This included upregulation of the casual gene (p<10^-3^), down-regulation of the paralog (p<10^-3^), or both (p<0.05). We repeated this analysis for genes with multiple paralogs by combining CGP pairs of the same causal gene (see Methods, Figure 4D). The frequency of each scenario in the combined dataset including all genes was significantly larger than expected by chance (Figure 4F). This suggests that imbalance in general, and specifically imbalance due to paralog down-regulation, occurs preferentially in the disease tissue.

An example for down-regulation of a paralog is presented by the charged multivesicular body protein 1a (CHMP1A) gene. CHMP1A is causal for pontocerebellar hypoplasia type 8, an autosomal recessive neurodevelopmental disorder [27]. Two germline mutations in CHMP1A were identified independently in patients, both leading to lack of CHMP1A expression in patient-derived cells [28]. Our data shows ubiquitous expression of CHMP1A across tissues, with intermediary expression in the cerebellum (Figure 5A, left). CHMP1A has a paralog, CHMP1B, with considerable sequence identity (55.6%) and significantly overlapping protein interaction partners. CHMP1B was significantly under-expressed in the cerebellum (Figure 5A middle), leading to imbalance and potentially insufficient compensation specifically in the disease tissue (Figure 5A, right). Additional examples appear in Figure 5B-E.

**Figure 5.**
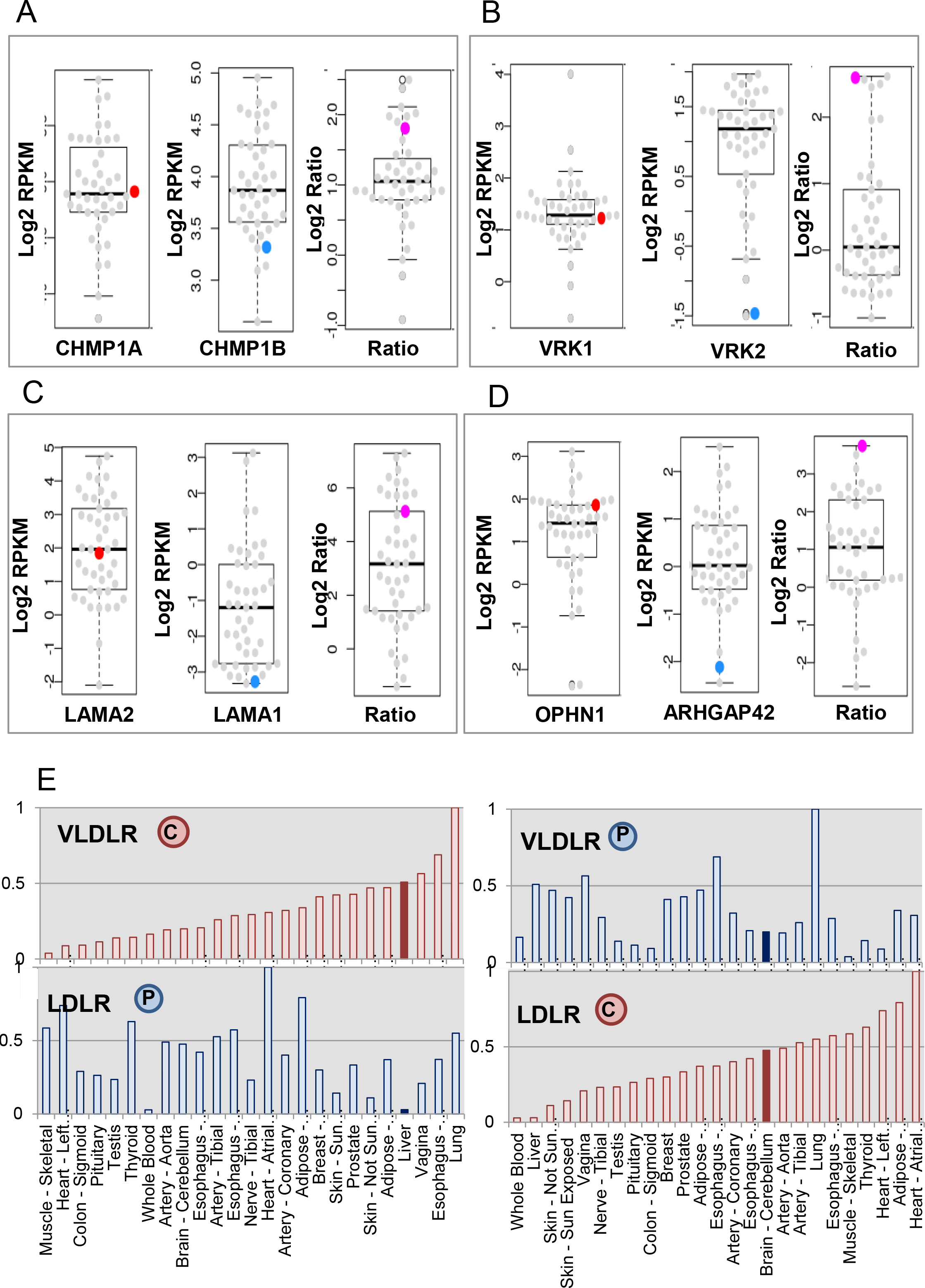
Specific examples for imbalanced expression of causal genes and their paralogs occurring preferentially in the disease tissues. Each panel shows the expression levels of the causal gene (left), the paralog (middle) and their ratio (right) in every tissue. Colored points indicate the respective values in the disease tissue. In each example the paralog of the causal gene is under-expressed in the disease tissue. A. CHMP1A and CHMP1B are two paralogous members of the CHMP family of proteins, with 55.6% sequence identity and significantly overlapping interactions. CHMP1A is causal for pontocerebellar hypoplasia type 8 that manifests in the brain, where CHMP1B is down-regulated (log_2_FC=-1, p=2.47*10^-4^). B. VRK1 and VRK2 are two paralogous members of the vaccinia-related kinase family of serine/threonine protein kinases, with 45% sequence identity and significant co-expression. VRK1 is causal for pontocerebellar hypoplasia type 1A that manifests in the brain, where VRK1 is not up-regulated (log2FC=-0.42, p=4*10^-3^), but VRK2 is strongly down-regulated (log2FC=-2.5, p=1.3*10^-18^). C. LAMA2 and LAMA1 are two paralogous laminin alpha subunits with 45% sequence identity. LAMA2 is causal for congenital muscular dystrophy that manifests in skeletal muscle. LAMA2 is over-expressed in muscle (log2FC=0.93, p=4.4*10^-10^), whereas LAMA1 is very lowly expressed (log2FC=-2.47, p=2.86*10^-9^). D. OPHN1 and ARHGAP42 are two paralogous Rho-GTPase-activating proteins with 51% sequence identity. OPHN1 is causal for X-linked mental retardation with cerebellar hypoplasia that manifests in the brain, where ARHGAP42 is strongly down-regulated (log2FC=-2.9, p=1.04*10^-12^). E. LDLR and VLDLR are two paralogs that cause distinct tissue-selective diseases and are down-regulated at each other's disease tissue. LDLR is causal for familial hypercholesterolemia that manifests in the liver (left), where it is expressed at intermediate level, while its paralog, VLDLR (48% sequence identity and significant overlap in protein-protein interactions), is down-regulated (log2FC=-2.5, p=1.32*10^-9^). VLDLR, in turn, is causal for cerebellar hypoplasia and mental retardation. VLDLR is expressed at intermediate level in the cerebellum (right), where LDLR is significantly under expressed (log2FC=-0.86, p=-0.02).

## DISCUSSION

Paralogous compensation is a key mechanism for maintaining genetic robustness [12–15,25]. Paralogs result from gene duplication events, and may be retained in the genome following sub- or neo-functionalization [29] or the sharing of gene dosage [30]. Their ability to compensate for each other relies on their functional similarity, and in some cases involves changes in the abundance of the functional paralog, its cellular localization or protein interactions (reviewed in [11]). Paralogous compensation is more frequent among young paralogs that have not diverged much [15,25], but was also observed among ancient paralogs [31]. Here we harnessed the concept of paralogous compensation to illuminate a fundamental question: What makes certain cell types or tissues succumb to a germline aberration while others remain robust. We hypothesized that paralogous compensation acts in various tissues throughout the body, however is limited and thus insufficient in the disease tissue, which therefore becomes vulnerable to germline mutations (Figure 1).

We tested our hypothesis on genes causal for 112 hereditary diseases that manifest predominantly in a single tissue (Figure 2). The causal variants of the genes we analyzed contained various types of aberrations, several of which can lead to partial or complete loss of the gene product or its function, due to, e.g., protein truncation or inframe missense mutations [23,24]. Other aberrations can lead to gain-of-function for which paralogous compensation may not be relevant, but these were shown in a systematic screen to occur at low frequency [23]. The causal genes and their paralogs in our dataset appeared to be functionally related by various measures, and, in accordance, were less essential than singleton genes (Figure 2D,E).

Dosage sharing between paralogs was suggested to be one of the first steps following gene duplication [30]. Here, we analyzed systematically for the first time dosage relationships between genes that are causal for tissue-selective hereditary diseases and their paralogs. For this, we exploited transcriptional profiles of 45 human tissues sampled from deceased donors with no genetic diseases [22]. Together, these profiles provided an atlas of gene expression in normal tissues and a baseline indicating the relevance of a gene within a specific tissue. While expression levels of wildtype alleles in patients carrying causal aberrations might differ from the levels observed in the general population, such changes were assessed previously and shown to be very limited [32–34]. Using these data, we found that the ratio between the expression levels of causal genes and their paralogs tend to be significantly high particularly in their disease tissues (Figure 3). This agrees with our hypothesis that, upon causal aberration, paralogous compensation will be limited specifically in the disease tissue, thereby making this tissue more vulnerable than other tissues expressing the same causal gene.

The relatively high expression ratios could stem from up-regulation of the causal gene in the vulnerable tissue, or from down-regulation of its paralog (Figure 1). To differentiate between these scenarios, we carried a rigorous differential expression analysis, which was enabled by the large numbers of samples available per tissue. The largest fraction of the cases included causal genes that were up-regulated significantly in their disease tissues, as previously observed [5,6]. However, in many other cases the causal gene was not upregulated and there was no tissue-specific isoform, yet a paralog was down-regulated significantly (Figure 4E and Figure 5). Notably, the frequency of each of these scenarios was higher than expected by chance (p<0.05, randomization test). A specifically interesting example involved two paralogous causal genes that were down-regulated at each other's disease tissue (Figure 4E). LDLR and VLDLR belong to the low density lipoprotein receptor gene family and are functionally overlapping (>47% sequence identity and significantly overlapping interactions). LDLR is causal for familial hypercholesterolemia that manifests in the liver, and VLDLR is causal for cerebellar hypoplasia and mental retardation. Each of them was expressed at intermediate levels at its respective disease tissue, and down-regulated at the disease tissue of its paralog, suggesting that by this they elicit tissue-specific phenotypes.

There remains a subset of causal genes for which imbalanced expression or significant expression changes were not observed. This includes cases where paralogous compensation may be irrelevant, due to limited functional overlap between paralogs or their tissue-specific isoforms, or a gain-of-function aberration. Interestingly a recent study showed that some paralogs in yeast are dependent on each other, and thus when mutated impart fragility rather than robustness [34]. We identified one such disease-related pair in human. BRAF and RAF1 are two functionally-related paralogous genes of the RAF family of serine/threonine protein kinases, known to be causal for various types of Noonan and Leopard syndromes. Interestingly, they were shown to form a heterodimer, thus explaining their dependency and common phenotypes [35]. Additional cases of paralogous compensation may remain hidden due to under-sampling of relevant cell types, or to lack of post-transcriptional profiling [33]. These might come to light upon analyzing data from specific cell types, or by rigorous proteomic analyses at large scales. In the future, it will be intriguing to extend the concept of compensation to higher-order entities such as pathways (e.g., [15,36]).

Our results show that systematic analysis of large-scale datasets illuminates dosage relationships and paralogous compensation events. They suggest that compensatory factors underlie tissue-selective genotype-phenotype relationships and particularly disease susceptibility, and point to paralogs as new and effective modifiers of tissue robustness.

## METHODS

### Disease set and genes analyzed in this study

The disease set included hereditary diseases with known protein-coding causal genes according to OMIM [37] that were predicted to manifest in either the brain, heart, liver, skeletal muscle, skin, testis or thyroid [6]. We used literature and expert curation to validate their clinical association and to filter out diseases with multiple affected tissues. Causal genes were downloaded from OMIM [37]. Paralogs were extracted from Ensembl-Biomart [6,38,39] and limited to paralogs with reciprocal sequence identity of 30% or more.

### Gene expression analyses

RNA sequencing profiles were obtained from the GTEx portal on 2/22/17 (version 6p) [40]. Only samples from individuals with traumatic injury as cause of death were included as proxy for healthy tissues. We verified the absence of possible confounders, including gender and age group, by applying mixed linear models to each tissue separately. To evaluate the expression distribution of a gene across tissues, we considered a gene as expressed in a certain tissue if its level exceeded 0.3 RPKM in at least half of the samples of that tissue. For each gene, the number of tissues expressing that gene was recoded. We included in the analyses only CGP pairs that were co-expressed in at least 5 tissues. Ratios between RPKM gene expression levels were calculated per sample. A distinct ratio was computed for a causal gene and each of its paralogs. In the combined analysis, a ratio was computed between the expression level of a causal gene and the sum of the expression levels of its paralogs. The ratio in a specific tissue was set to the median ratio across all samples of that tissue.

### Functional overlap analysis

We correlated between the expression levels of a causal gene and each of its paralogs across tissues by using Pearson correlation. For each gene, its expression level per tissue was set to the median RPKM level over samples of that tissue. We downloaded data of experimentally-detected protein-protein interactions from BioGRID [41], DIP [42] and IntAct [43] by using myProteinNet [44] and computed the number of proteins that interact with a causal gene, with its paralog, and with both. The CRISPR scores of genes, which represent their essentiality, were extracted from [16].

### Differential expression analysis

Differential expression analysis was applied to 37 GTEx tissues with at least five samples. Raw counts were extracted from GTEx portal and normalized using the TMM method by the edgeR package (27), to obtain the same library size for every sample. Genes with less than 10 counts in all samples were removed before normalization. In each sample, we transformed RNA-sequencing normalized counts using VOOM [45], and calculated differential expression using a linear model in the R-package Limma [46]. Specifically, all samples of the same tissue were compared to all other samples. Only genes with an absolute 2-fold change and FDR adjusted P-values <0.01 were considered differentially expressed.

### Statistical analysis

We used mixed linear models to predict the expression of a causal gene in each tissue by the expression levels of its paralogs, by the number of its paralogs, by whether its expression was measured in the disease tissue and by the amount of tissues which manifest a disease associated with this gene. We accounted for the clustered structure of the donors by including a random intercept in all of the models. Mixed linear models were computed by using IBM SPSS Statistics, Version 23.0. We compared between the expression distributions of causal genes and their paralogs across tissues by using the Kolmogorov-Smirnov test. The significance of protein interaction overlap between a causal gene and its paralog was computed by using Fisher exact test. We compared between essentiality scores of causal genes and protein-coding genes without paralogs by using the Kolmogorov-Smirnov test. We used a randomization test to assess whether causal genes (and their paralogs) are overexpressed (or under-expressed) preferentially in the disease tissue relative to other tissues. Specifically, in each randomized run each causal gene was assigned a randomly selected disease tissue out of the set of GTEx tissues expressing the causal gene and its paralog. For causal genes with multiple paralogs, the disease tissue was selected randomly from the set of GTEx tissues expressing the causal gene and at least one of its paralogs. We then counted the number of causal genes that were significantly over-expressed, had an under-expressed paralog, or both, in the randomly selected disease tissues. We repeated this analysis 1,000 times. Statistical significance was set to the fraction of randomized runs in which the number of causal genes in a given subset was at least as high as the fraction observed for these pairs in the original dataset.

## ACKNOWLEDGEMENT

We thank Francois Aguet, Ayellet Segre and Vered Chalifa-Caspi for their help in the gene expression analyses of GTEx samples. We thank Liad Alfandri for his help in the association between diseases and tissues.

